# A DNA mass conservation mechanism underpins cellular mtDNA number regulation

**DOI:** 10.64898/2026.07.08.737183

**Authors:** Jagir R. Hussan, Asgeir Kobro-Flatmoen, Peter Ruoff, Stig W. Omholt

## Abstract

The nucleoid, which houses mtDNA within the mitochondrial matrix, is a phase-separation-driven biomolecular condensate capable of carrying out a broad spectrum of complex functions, including DNA replication, transcription, and repair. Here, we show by data-driven computational modelling that the concept of a tightly regulated intranucleoid de-oxynucleoside triphoshate (dNTP) pool explains the observation that the number of mtDNA base pairs per cell is conserved in human hybrid cell lines regardless of the size of the introduced mitochondrial genome. This concept is then used to address the enigmatic observation that the synthesis rate of the short DNA strand called 7S DNA, which is part of the triple-stranded displacement loop (D-loop) found in the main noncoding region of mtDNA, increases dramatically during the cell cycle. Collectively, our quantitative analyses suggest that the mammalian mtDNA replisome uses a strictly controlled intranucleoid dNTP pool based predominantly on the synthesis and degradation of 7S DNA. One potential evolutionary explanation for this mechanism is that it offers an energetic advantage by enabling greater reliance on the salvage pathway for mtDNA replication.

## Introduction

Although there is a ∼ 50-fold variation in median cellular mtDNA number among human mitotic and postmitotic tissues (Bogenhagen, 2012; Rath et al., 2024), the variation in the mtDNA number between cells taken from a clonal population is quite moderate (O’Hara et al., 2019). Therefore, some kind of regulatory control needs to be involved. Despite the extensive attention mitochondrial biology has received over several decades, revealing that mitochondria perform a dauntingly complex set of bioenergetic, metabolic and signaling functions in eukaryotic cells (Monzel et al., 2023), we still do not grasp how this regulation occurs.

In most eukaryotic cells, mitochondria are dynamic organelles that fuse and divide to form constantly changing tubular networks (Westermann, 2002). Mitochondrial DNA is usually located in nucleoprotein complexes called nucleoids that can move within this network (Alán et al., 2016). An ultrastructure study of nucleoids in a L1210 leukemia cell line reported that the nucleoid core is generally surrounded by a thick electron-lucent layer, presumably subdivided by walls into tens of compartments (Prachař, 2016). The cellular number of nucleoids varies from several hundreds to thousands (Legros et al., 2004; Bogenhagen, 2012; Ježek et al., 2019), with reported sizes spanning 31 to 318 nm (Brown et al., 2011; Kopek et al., 2012). Nucleoids are organised into higher-order assemblies, including respiratory chain supercomplexes and endoplasmic reticulum–mitochondria complexes (Lee and Han, 2017). Beyond acting as a protective shield for mtDNA, more than 50 nucleoid-associated proteins contribute to mtDNA maintenance and gene expression by forming transient or stable associations with mtDNA itself or with other nucleoid-associated proteins (Lee and Han, 2017). The number of wild-type (WT) mtDNAs contained in one nucleoid has been a controversial issue for years. The estimates of the mean number range from one to ten (Satoh and Kuroiwa, 1991; Legros et al., 2004; Bogenhagen, 2012; Kukat et al., 2011; Pavluch et al., 2023), and, at least in human cell cultures, the intranucleoid variation around the mean appears to be quite restricted (Legros et al., 2004).

The elegant experiments of Tang *et al*. (Tang et al., 2000) showed that when human cells devoid of endogenous mtDNA were supplied with mitochondria from Kearns–Sayre syndrome patients containing either exclusively wild-type mtDNA (16,569 bp), exclusively partially duplicated mtDNA (25,323 bp), or exclusively partially deleted mtDNA (9,049 bp), the total number of mtDNA base pairs per cell was essentially the same across all three cybrid lines (a phenomenon hereafter referred to as mtDNA mass conservation). The authors suggested that mtDNA mass could be conserved if the mtDNA number was regulated by tightly controlled mitochondrial dNTP pools (Tang et al., 2000), but did not elaborate on this idea in terms of how such control could be achieved.

The existence of a highly dynamic tubular network (Westermann, 2002) makes the concept of a distinct mitochondrion and mitochondrial matrix quite evanescent. Thus, it is challenging to contemplate how the mitochondrial matrix *per se* could be the location of a control module capable of providing a distinct number of dNTPs to the replication machinery in each nucleoid. However, the fusion and fission dynamics is of no concern if we assume that each nucleoid is equipped with a unique and tightly regulated dNTP pool. Since a nucleoid does not have a membrane (Lee and Han, 2017), the contention of an intranucleoid dNTP pool may seem questionable. However, the fact that nucleoids move within the mitochondrial matrix (Alán et al., 2016) clearly supports the notion that they represent coherent structures capable of shielding their cargos from the environment of the mitochondrial matrix. This notion is substantiated by the demonstration that mitochondrial transcription factor A (TFAM) undergoes liquid-liquid phase separation with mtDNA to drive nucleoid self-assembly and that nucleoids act as membrane-less organelles into which mitochondrial transcription complexes can be incorporated and promote the retention of nucleoside triphosphates (Long et al., 2021). Thus, nucleoids appear to be a firm member of the rapidly expanding class of phase-separation-driven biomolecular condensates that can sport a wide range of functions facilitated by a complex internal organisation (Alberti, 2017). In fact, the three main functions that we currently attribute to the nucleoid based on extensive experimental data—replication, transcription, and repair of mtDNA (Gustafsson et al., 2016; Lee and Han, 2017; Kang et al., 2018)—represent a par excellence demonstration of the bewildering regulatory complexity that can be embedded in a phase-separation-driven biomolecular condensate. In this context, the existence of a tightly regulated intranucleoid dNTP pool does not appear implausible.

The major non-coding region of human mtDNA spans approximately 1.1 kb and a large part of this region often incorporates a linear third DNA strand (called 7S DNA) of approximately 650 nucleotides (nt), forming a stable D-loop structure (Nicholls and Minczuk, 2014). The half-life of 7S DNA is about 70 minutes (Bogenhagen and Clayton, 1978) and about 95% of all replication events terminate to produce 7S DNA only (Falkenberg and Gustafsson, 2020). Since its discovery more than 50 years ago, the function of the D-loop has remained enigmatic (Nicholls and Minczuk, 2014). However, more than 15 years ago Antes *et al*. demonstrated that in HeLa cells the ratio between the cellular number of 7S DNA and the number of mtDNAs in the S phase is approximately seven times higher than at the G1/S boundary (Antes et al., 2010). This observation led the authors to suggest that the mitochondrial nucleotide pool can be significantly influenced by 7S DNA degradation (Antes et al., 2010), but they did not provide any supporting quantitative analysis or elaborate on the reasons why a cell should synthesise 7S DNA to provide the mitochondrial matrix with nucleotides. Intriguingly, the concept of an intranucleoid dNTP pool opens up the possibility that 7S DNA based dNTP production and utilisation is a pure intranucleoid operation.

In the following, using a continuous/discrete event (hybrid) model within an object-oriented framework, we first identify control principles for the dynamics of an intranucleoid dNTP pool that would lead to the conservation of mtDNA mass in cybrid non-dividing cell lines. Within the same computational framework, using the time-resolved 7S DNA production data reported by Antes *et al*. (Antes et al., 2010) and assuming that 7S DNA is the primary source of dNTPs for mtDNA replication, we subsequently demonstrate that the observed temporal increase in nucleoid number from early G1 to G2/M in HeLa cells (Sasaki et al., 2017) can be reproduced. On the basis of these cell cycle results, it is shown that the cybrid mtDNA mass conservation results reported by Tang *et al*. (Tang et al., 2000) can be closely reproduced. We also provide a possible evolutionary reason based on energetic considerations for the existence of this counterintuitive cellular strategy.

Taken together, our findings support the conception that the dNTPs used for mtDNA replication are predominantly derived from 7S DNA produced within the nucleoid and that the number of dNTPs provided for each replication event is tightly regulated. This offers a fresh perspective that may help in resolving key questions currently facing mitochondrial regulatory biology.

### Conservation of mtDNA mass in a non-dividing cell

Tang *et al*. observed mtDNA mass conservation in mitotic 143B206 cell lines (Tang et al., 2000), and to our knowledge, no similar study has been conducted on non-dividing cells. However, if there is a mechanism that causes this conservation in mitotic cells in general, it is arguably also present in quiescent mitotic cells and postmitotic cells. For simplicity, we therefore used the non-dividing cell case to evaluate to which degree an intranucleoid dNTP pool concept would lead to mtDNA mass conservation under a range of conceivable regulatory conditions. To this end, we constructed a hybrid model in which nucleoids are entities (objects) that combine data (attributes) and behaviour (methods) that operate autonomously except that the dNTP pool filling rate is under cellular control. Since the number of nucleoids (or mtDNA) at a given time point emerges from the internal logic of the computational model, there is no need to seek an explicit equation describing the temporal dynamics of nucleoids and mtDNAs. The premises of the model are presented below. The Python code needed to produce all results reported in this article is available at https://github.com/Jagirhussan/mtDNA-mass-conservation. The code was deliberately designed to facilitate the inclusion of much more biological detail than we have deemed necessary for the purposes of this study.

Assuming that the nucleoid number in a non-dividing cell in a fixed environment fluctuates around an approximately stable mean value, we let the cell regulate the number of nucleoids by means of a negative feedback controller. The controller is assumed to sense directly or by proxy, the number of nucleoids relative to its set point and use this information to regulate the supply rate of dNTPs to all intranucleoid pools (*r(t)*). Phenomenologically, this regulation can be described by the function

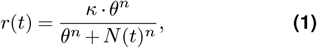

where *κ, θ* and *n* are control parameters and *N(t)* is the number of nucleoids at time *t*. The function expresses that the dNTP supply rate increases when the number of nucleoids is below the target value and decreases when it is above. For the purposes of this study, it is not necessary to further detail the regulatory anatomy of the controller. Nevertheless, it is worth pointing out that it is likely to be quite intricate, considering that the target value of the nucleoid number will have to be a function of numerous parameters by which the cellular condition is monitored.

Initiation of mtDNA replication is supposed to be under the control of the regulatory machinery within the nucleoid and to start when the intranucleoid number of dNTPs exceeds a given threshold. The cybrid line data reported by Tang *et al*. are consistent with the notion that the mechanism responsible for determining the threshold size is oblivious to changes in mtDNA genome size. We therefore assumed that the scaling of the threshold size is under cellular control and that this scaling defines the mean number of wild-type intranucleoid mtDNAs specific to a given cell type (Legros et al., 2004).

Tang *et al*. (Tang et al., 2000) did not measure the number of nucleoids in their cybrid cell lines. Based on the reported wild-type cellular mtDNA number and assuming that each nucleoid contained five mtDNAs, which is well within the range observed in different human cell cultures (Legros et al., 2004; Bogenhagen, 2012; Pavluch et al., 2023), the mean cellular number of nucleoids would have been 2,611. To allow a direct comparison between the model predictions and the data reported by Tang *et al*., we let the non-dividing wild-type cybrid cell contain approximately the same mean number of nucleoids.

We assumed that the replication time of all mtDNAs within each nucleoid was one hour (Bogenhagen and Clayton, 1978; Sasaki et al., 2017), and let, for convenience, the hybrid model have a time resolution of one hour.

Due to the paucity of constraining data, we allowed for a normal-distributed nucleoid lifespan (*N* (*µ, CV* · *µ*)) and a comparable random mortality of nucleoids in both wild-type and mutants. In line with data on mtDNA turnover rate in quiescent rat hepatocytes (Kai et al., 2006), we let *µ* = 100 h. The coefficient of variation (CV) was set to 0.2. Furthermore, in the two mutant cases (9,049 bp and 25,323 bp), we allowed the dNTPs that remained after a replication event to be lost or retained. If retained, the dNTPs could remain in the mother nucleoid or be transferred to the daughter nucleoid during nucleoid division. Finally, the mechanism controlling the filling of the dNTP pool could either account for the dNTPs remaining after a preceding replication event or it could be oblivious to the number of remaining dNTPs and fill the pool with a fixed number of dNTPs each time. This set of options left us with 10 different regulatory scenarios for each of the mutants and two wild-type scenarios, since in the latter case we assumed for simplicity that there were no dNTPs left after a replication. In every scenario, we assumed that mtDNA replication proceeds until the available dNTPs are insufficient to support another round of replication. At that point, the nucleoid rapidly divides, randomly distributing both the pre-existing and newly synthesized mtDNAs between the mother and daughter nucleoids.

Based on the above premises and regulatory options, we let the hybrid model predict the temporal dynamics of nucleoids and mtDNAs in a single non-dividing cybrid cell for the 22 different cases enlisted. In the wild-type, after a transient run-in period, the model expectedly predicts that the number of nucleoids fluctuates stably around the targeted mean value (Fig. 1A). The figure shows the case where the nucleoids have a normal-distributed lifespan, but the use of a constant hourly random mortality rate gave a similar temporal pattern. If the number of nucleoids in a non-dividing cell is abruptly perturbed upward or downward, the hybrid model predicts that the mean value of nucleoids moves back to essentially the original pre-perturbation value (Fig. 1A), i.e. it shows almost perfect adaptation. In line with this, cell cultures exposed to severe depletion of their mtDNA population by ethidium bromide (EtBr) recover their original mtDNA numbers (and thus nucleoid numbers) in 10-14 days (Kristiansen et al., 2023). The height and width of the amplitude will depend on several factors such as the parameterisation of Eq. (1), the type of initialisation regime used, or the type of negative feedback controller used. However, since only the mean steady state number of nucleoids was used to assess mtDNA mass conservation, we did not conduct a systematic analysis of this variability.

**Figure 1.**
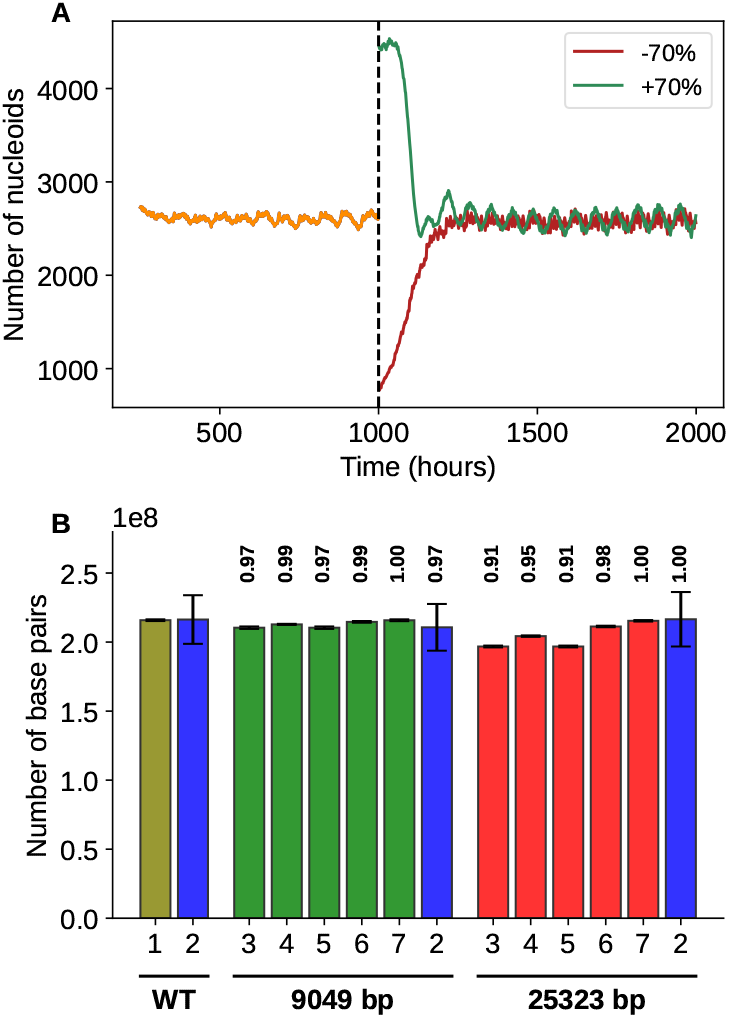
Predicted nucleoid dynamics and mtDNA mass conservation in a non-dividing cell with wild-type mtDNA. A. 250 < *t* < 1000: Predicted temporal variation in the number of nucleoids (N(t)) after a transient run-in period (a single run). *t* ≥ 1000: The predicted effect of abruptly increasing or decreasing N(t) by 70% at t=1000. B. Predicted cellular number of mtDNA base pairs in the three cybrid cases versus experimental values obtained by (Tang et al., 2000). 1: WT; 2: Experimental values for cybrid cells with WT mtDNA, severely depleted mtDNA (9,049 bp) and partially duplicated mtDNA (25,323 bp), respectively; 3: Excess dNTPs are lost between each replication; 4: Excess dNTPs retained in the mother nucleoid; 5: Excess dNTPs retained in the daughter nucleoid; 6: Excess dNTPs retained in the daughter nucleoid are accounted for during refilling of the dNTP pool; 7: Excess dNTPs retained in the daughter nucleoid do not affect the refilling of the dNTP pool. Each of the 11 predicted cases shows the grand mean of 50 model runs, and the error bars represent the third standard deviation to allow visibility. Each value shown above the green and red bars represents the ratio of the predicted mean number of base pairs in the mutant cybrid cell to the predicted mean number of base pairs in the wild-type cybrid cell. The values above the two blue bars are the base pair ratios mutant to wild-type reported by Tang *et al*. (Tang et al., 2000). Parameter values: *κ*=4,420, *θ*=3,000, n=2, *µ*=100, CV=0.2. Each numerical experiment was initiated by constructing a nucleoid pool (N = 2,611), where each nucleoid was attributed a lifespan drawn from the above normal-distribution, and a dNTP pool size that was randomly drawn from the interval [0, 165690 (16569 ·2·5)].

The model predicts that in a cybrid cell containing 9,049 bp mtDNA, the presence of an intranucleoid dNTP pool results in nearly complete preservation of the mtDNA mass compared to the wild type regardless of the option by which surplus dNTPs are processed (Fig. 1B). In a cybrid cell containing 25,323 bp mtDNA, the picture is more nuanced and perfect conservation is only achieved if excess dNTPs are retained in the daughter nucleoid and the pool filling mechanism is oblivious to the number of dNTPs remaining after the previous replication (Fig. 1B). Almost identical results were obtained using a constant hourly mortality rate instead of a normal-distributed lifespan (figure available at https://github.com/Jagirhussan/mtDNA-mass-conservation).

It should be emphasised that for any targeted mean number of nucleoids, the manifestation of mtDNA mass conservation is insensitive to the mean number of mtDNAs in each nucleoid, the mtDNA replication time, and the mean lifespan of the nucleoids, as any change in these three variables can be accommodated by adjusting the parameters in Eq. (1). Thus, within our computational framework, the concept of a tightly controlled intranucleoid dNTP pool robustly leads to the same mean number of mtDNA base pairs in non-dividing cells that are exposed to the same environmental conditions and only differ by their mtDNA genome size.

### 7S DNA based nucleoid replication during the cell cycle

The increase in the number of nucleoids follows a power function throughout most of the cell cycle in HeLa cells (Sasaki et al., 2017). Given the existence of intranucleoid dNTP pools, this indicates that the dNTP supply rate to the pools also follows a power function since the replication rate of nucleoids is a function of how fast their pools fill up. Intriguingly, in HeLa cells, the increase in the cellular level of 7S DNA appears to also follow a power function throughout the main part of the cell cycle (Antes et al., 2010). And RNAi knockdown of the 7S DNA-protective mitochondrial single-stranded DNA-binding protein (mtSSB) in HeLa cells causes, within three days, a more than 80% drop in the number of 7S DNAs and a concomitant 30% decrease in the number of mtDNAs due to a reduction in the rate of mtDNA synthesis (Ruhanen et al., 2010). These findings motivated us to examine the radical hypothesis that close to all dNTPs delivered to intranucleoid dNTP pools needed to double the nucleoid population before cell division originates from 7S DNA. In this scenario, dNTPs will have to be imported into the nucleoid from the mitochondrial matrix and used to synthesise 7S DNA, which within the nucleoid is then degraded by an ssDNA nuclease such as mitochondrial genome maintenance exonuclease 1 (MGME1) (Kornblum et al., 2013; Szczesny et al., 2013; Nicholls et al., 2014; Yang et al., 2018; Mao et al., 2024) into deoxynucleoside monophosphates (dNMPs). Subsequently, these dNMPs are phosphorylated by nucleoside monophosphate kinases (NMPKs) and nucleoside diphosphate kinases (NDPKs) (Van Rompay et al., 2000; Xu et al., 2008; Blanco and Blanco, 2017) to regenerate dNTPs, which are then retained within the nucleoid.

To estimate the temporal development of the cellular number of 7S DNA during the cell cycle we used the observation that the ratio between the cellular number of 7S DNA and the number of mtDNAs in the S phase is approximately seven times higher than at the G1/S boundary (Antes et al., 2010). We assumed that the fold change (*F* (*t*)) in the cellular number of 7S DNA from early G1 (EG1) to late S (LS) could be approximated by the power function:

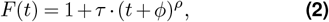

premised on that there is a monotonic temporal increase in the cellular number of 7S DNA and that it is > 0 at the beginning of the cell cycle. The cellular number of 7S DNA throughout the cell cycle (7*SDNA*(*t*)) is then given by:

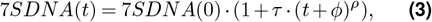

where 7SDNA(0) is the cellular number of 7S DNA at the beginning of EG1. We estimated the parameters *τ, ϕ* and *ρ* by using a lowest mean square error (MSE) method for the initial 7S DNA values in the set {7SDNA(0) ∈ ℤ: 500 ≤ 7SDNA(0) ≤ 3000 and 7SDNA(0)/500 ∈ ℤ}. In each case, we assumed that the cellular number of nucleoids (N(t)) (each containing 5 mtDNAs) in EG1 was N(0) = 490 (Sasaki et al., 2017), in the set {N(7) ∈ ℤ: 510 ≤ N(7) ≤ 650 and N(7)/10 ∈ ℤ} at G1/S (Sasaki et al., 2017) and in the set {N(13) ∈ ℤ: 680 ≤ N(13) ≤ 990 and N(13)/10 ∈ ℤ} in S (Sasaki et al., 2017), and we demanded that 7SDNA(7)/5 ·N(7) = 2.6 and 7SDNA(13)/5· N(13) = 18.3 to ensure compliance with the data reported by Antes *et al*. (Antes et al., 2010). From the resulting family of 7SDNA(t) curves we selected the one with the lowest MSE (Fig. 2A) and the associated values of *τ, ϕ*, and *ρ* were recorded for further use.

**Figure 2.**
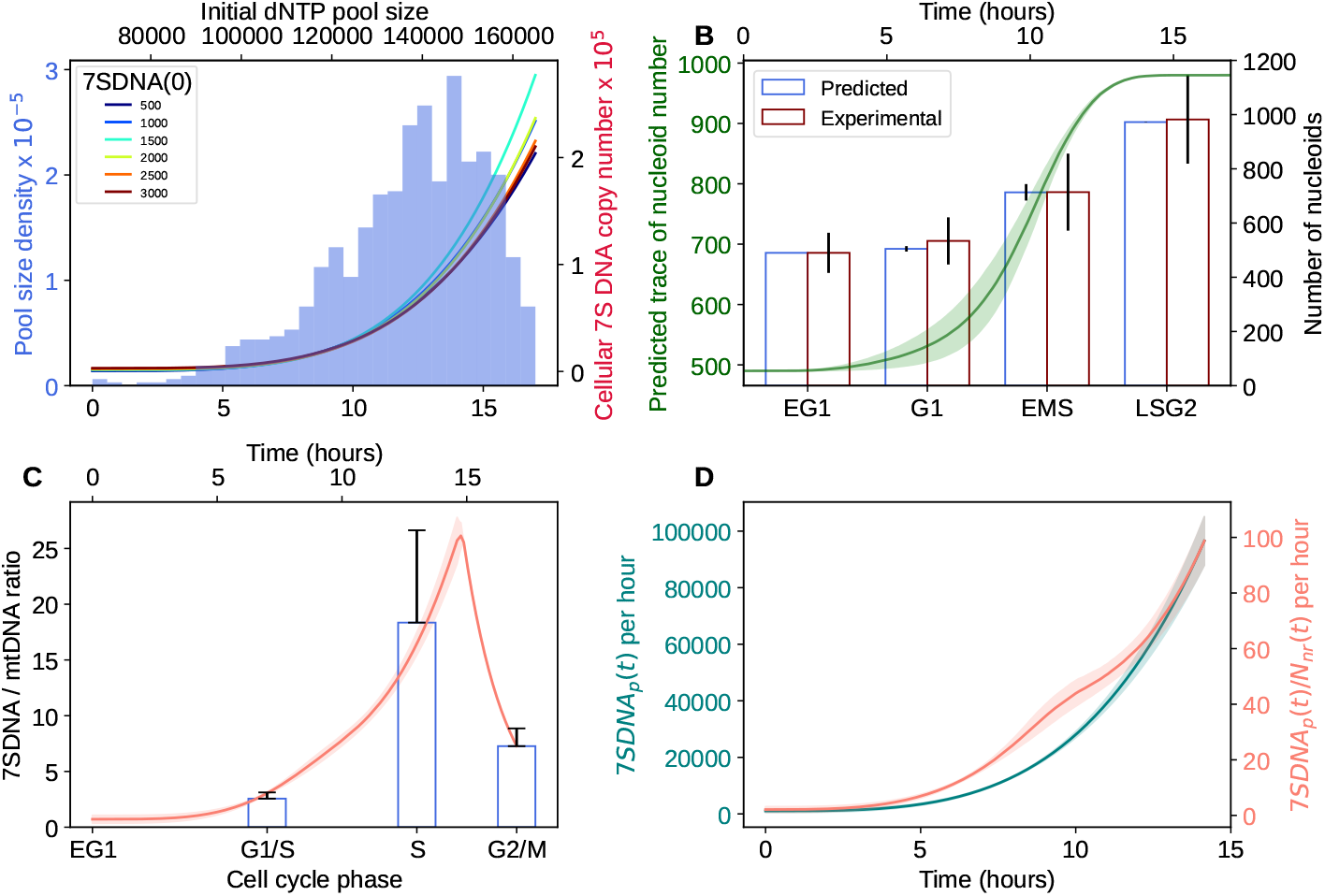
Nucleoid and 7S DNA dynamics throughout the cell cycle. A. Top x-axis and left y-axis: Example of an initial dNTP pool size distribution based on a discrete beta-binomial distribution with *n* = 165,690, *α* = 10 and *β* = 2. Bottom x-axis and right y-axis: Estimated cellular 7S DNA number throughout the cell cycle until end of G2 (*t* = 17) as a function of the initial number of 7S DNA at *t* = 0 (7SDNA(0)) based on Eqs. (2) and (3). B. Top x-axis and left y-axis: Predicted number of nucleoids from the beginning of EG1 (*t* = 0) to end of G2 (*t* = 17) when we assume that replication of nucleoids is inhibited shortly after duplication of the initial population of nucleoids. The graph shows the mean±SD when using all the six 7SDNA(0) values depicted in panel A. Bottom x-axis and right y-axis: Predicted (mean±SD) vs reported (Sasaki et al., 2017) number of nucleoids in a HeLa cell in four cell cycle phases. EG1, early G1 phase; EMS, early S/mid S phase; LSG2, late S/G2 phase. C. Predicted vs reported (Antes et al., 2010) cellular 7S DNA number relative to the number of mtDNAs. The graph shows the mean SD when using all the six 7SDNA(0) values depicted in panel A. D. Left y-axis: Estimated cellular production rate of 7S DNA (mean ± SD) from the beginning of EG1 to *t* = *t*_*d*_ for all six 7SDNA(0) values depicted in panel A. Right y-axis: The predicted production rate in each non-replicating nucleoid (*N*_*nr*_ (*t*)) (mean ± SD) for the same time period and same 7SDNA(0) values. *N*_*nr*_ (*t*) for all six 7SDNA(0) values are depicted in Fig. S7.

Equation (3) was then used to obtain a differential equation that describes the time rate of change of dNMP throughout the cell cycle:

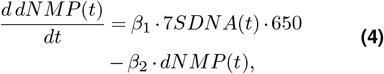

where *β*_1_ is the proportion of the 7S DNA population degraded to dNMPs per hour by an ssDNA nuclease and *β*_2_ · *dNMP* is the number of dNMPs equipped with two phosphate groups per hour by NMPKs and NDPKs. The cellular production rate of dNTPs as a function of time (*dNTP*_*rate*_(*t*)) is thus given by:

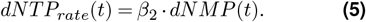

Assuming a half-life of 7S DNA of 70 minutes (Bogenhagen and Clayton, 1978), we let *β*_1_ = 0.592, and assuming a very fast phosphorylation of dNMPs we let *β*_2_ = 0.95.

To define the size of the dNTP pool in each nucleoid in the initial nucleoid population at the beginning of EG1, we sampled from a discrete beta-binomial distribution *X* ∼ BetaBin(n,*α,β*), where n is the dNTP pool size (165690 nt) at which replication starts and *α* and *β* are parameters determining the shape of the associated beta distribution (illustrated in Fig. 2A). For each pair of *α* and *β* values, we sampled 50 distributions.

In the case of a non-dividing cell (Fig. 1), we let the number of nucleoids be homeostatically regulated through cellular negative feedback control of the dNTP production rate within each nucleoid (Eq. (1). There is clearly no homeostatic regulation of nucleoid number in actively dividing cells, but there still has to be a regulatory scheme ensuring that the nucleoid number doubles during the cell cycle. Instead of introducing a speculative explicit control function, we mimicked its presumed operation by dividing the celluar dNTP production rate *dNTP*_*rate*_(*t*) obtained from Eqs. (3), (4) and (5) by the number of nucleoids that did not actively replicate mtDNA at time *t* (*N*_*nr*_(*t*)). By this we obtained the fill rate of the dNTP pool in each nucleoid as a function of time. The time resolution of this mitotic version of the hybrid model was one minute. For simplicity, we assumed that none of the nucleoids died during the cell cycle.

When we run the model up to the end of G2 (*t* = 17), we obtained a reasonable fit with the data reported by Sasaki *et al*. (Sasaki et al., 2017) except for the late S/G2 phase for a fairly wide range of 7SDNA(0) values (Fig. S1). The reason for this discrepancy is that the number of nucleoids increases dramatically at the end of the late S phase and becomes unrealistically high at the end of G2 (Fig. S1). But before this explosion in numbers, there is a fairly wide time window where the nucleoid population plateaus at twice the number of the initial population size (Fig. S1). This plateau is caused by a time lag between the division of the nucleoid from the initial population with the lowest initial dNTP pool size and the second division of the nucleoid from the initial population with the highest initial dNTP pool size (and the concurrent division of its daughter nucleoid from its first division).

Based on the data (Sasaki et al., 2017), we therefore found it reasonable to assume that the regulatory scheme alluded to above includes a mechanism to detect when the nucleoid population has duplicated and a mechanism for stopping 7S DNA production and mtDNA replication shortly after this (Fig. S2). We implemented this assumption in the model and found that in this case we were able to recapitulate the Sasaki *et al*. data pretty well throughout the cell cycle (Figs. S3 and 2B).

After termination of 7S DNA synthesis, the subsequent decline in the cellular number of 7S DNA can be described by the differential equation:

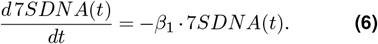

If 7S DNA synthesis is terminated shortly after duplication of the initial nucleoid pool at *t* = *t*_*d*_, it follows that the cellular number of 7S DNA relative to the mtDNA number at G2/M (*t* = 17) (*R*_*G*2*/M*_ ) can be estimated from the equation:

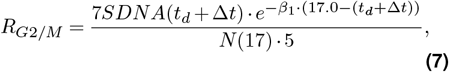

where N(17) is the number of nucleoids at the end of G2 and *t* +Δ*t* is the time point when 7S DNA synthesis stops and the 7S DNA population starts to decline. If we let Δ*t* = 0, the predicted 7S DNA/mtDNA ratio at G2/M is about 70% of the mean value reported by Antes *et al*. When Δ*t* ≈ 30 minutes, the predicted temporal development of the 7S DNA/mtDNA ratio from the beginning of EG1 to the end of G2 is fully consistent with the three reported data points (Antes et al., 2010) (Fig. 2C). This agreement supports the view that the 7S DNA synthesis rate increases monotonically from the beginning of the cell cycle to nearly the end of S phase, and that both mtDNA replication and 7S DNA synthesis are blocked during the remaining portion of the cycle through inhibition of an essential replisome component. It should be noted that very simple assumptions underpin the sharp peak of the graph, and possibly the peak is less distinct than what our model predicts. Antes *et al*. reported a substantial variation in the 7SDNA/mtDNA ratio in the S-phase (Fig. 2C). Considering that the S phase lasts ≈ 9 hours (Sasaki et al., 2017) during which the ratio increases dramatically (Fig. 2C), a large variability in the ratio is expected since it will depend on when in the S phase a synchronized cell population is sampled (Antes et al., 2010).

A 25% reduction or increase in the 7S DNA degradation constant *β*1 has a very moderate impact on the above results. The degradation rate of single strand DNA by MGME1 has been estimated to be 6.3 nt · s^-1^ in the 5’-to-3’ direction at 22^°^C by nuclease-induced stepwise photodropping (Chiu et al., 2024). Assuming that one MGME1 per nucleoid actively degrades 7S DNA, the cell will degrade ≈ 76,300 7S DNA in 2.2 hours (17.0 − (*t*_*d*_ + Δ*t*) = 17.0 − (14.3 + 0.5)) (*t*_*d*_ and Δ*t* are mean values for all six 7SDNA(0) cases). This gives a degradation constant value that is 67% of the one used. Considering that the rate of 7S DNA degradation by MGME1 is likely to be higher at 37^°^C and that the reported rate is in any case likely to be lower than the *in vivo* rate—since MGME1 cooperates with Pol *γ* and the mitochondrial twinkle mtDNA helicase in the degradation of linear mtDNA (Peeva et al., 2018)—this MGME1-based decay constant estimate is intriguingly consistent with the one we have used.

Since the use of more left-skewed initial beta-binomial distributions gave a poorer fit to the data (Sasaki et al., 2017) (Figs. S4 and S5), we predict that the distribution of the dNTP pool size at the beginning of the cell cycle in HeLa cells will be found to be rather right-skewed. Furthermore, based on how the dNTP pool size distribution develops from the time point when the initial nucleoid population is duplicated to the end of G2, we predict that the filling of the dNTP pools will be terminated in early G2 (Fig. S6). At this time point, the distribution is quite similar to the initial one, which will ensure that the temporal development in the number of nucleoids during the next cell cycle will be quite similar to the first (Fig. 2B).

Assuming that the cellular production rate of 7S DNA per hour (7*SDNA*_*p*_(*t*)) can be described by the equation:

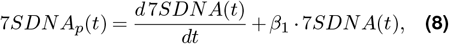

it follows from Eq. (3) that:

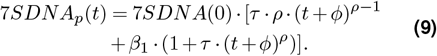

Using this equation, we calculated the maximum cellular and intranucleoid production rate per hour of 7S DNA (Fig. 2D). The results suggest that the maximum hourly intranucleoid production rate is ≈ 100, which requires that the replication time of a single 7S DNA must be less than 3 minutes in a nucleoid with five mtDNAs. Since this figure is far above the replication time that would be expected from the measured rate of nucleotide incorporation by Pol *γ* (Johnson and Johnson, 2001), it follows that as long as the number of Pol *γ* and other components of the mitochondrial replisome is not rate limiting, the nucleoid replication rate of 7S DNA is well below its maximum capacity even in late S. Although the predicted maximum 7S DNA synthesis rate is dependent on the assumed number of mtDNAs per nucleoid, a lower or a moderately higher number will not affect this conclusion.

In an asynchronous cell culture containing a large number of cells, the proportion of mtDNAs containing a D-loop (*D*_*prop*_) can be estimated from the equation:

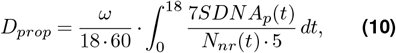

where *ω* is the synthesis plus residence time of a 7S DNA strand per mtDNA in minutes and 7*SDNA*_*p*_(*t*) = 0 for *t* > *t*_*d*_. If the Pol *γ* polymerisation rate is 37 nt per second (Johnson and Johnson, 2001), *ω* = 0.29. Assuming *ω* ∈ [0.29, 0.65], the predicted percentage range of D-loops is intriguingly concordant with the reported range of 5-12% in HeLa cells (Brown et al., 1978) (Fig. S7). This suggests a residence time approximately in the range 0 - 21 seconds. However, the observed variation may also have been influenced by variation in the synthesis time (Johnson and Johnson, 2001).

Together with the fact that the A, T, C, and G frequencies in 7S DNA are very similar to the respective frequencies in the complete human mitochondrial DNA sequence (Andrews et al., 1999) (7S DNA/whole genome): A:0.303/0.309; T: 0.242/0.247; C: 0.326/0.313; G: 0.129/0.131), the results support the contention that 7S DNA can be the main nucleotide source for mtDNA replication in general.

### mtDNA mass conservation with 7S DNA based dNTP supply

We then went back to the Tang *et al*. data (Tang et al., 2000) assuming that the dNTP supply to the intranucleoid dNTP pools also in this case had come solely from 7S DNA. We presumed that 143B206 cybrid cells had the same N(t) profile as the one depicted in Fig. 2B. The time axis of the profile was rescaled to account for a possibly slightly longer cell cycle time of ≈ 19.8 h (King and Attardi, 1989). Based on the reported mean cellular number of mtDNAs in the WT cybrid line (13,056) (Tang et al., 2000) and assuming five mtDNAs per nucleoid, we demanded 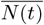 for the WT case to be ≈ 2,611, defining a target curve (Fig. 3A). To account for the increase in the number of nucleoids, we multiplied the righthand side Eq. (3) with the ratio of the mean number of nucleoids of the target curve from 0 to 19.8 h and the mean number of nucleoids of the curve depicted in Fig. 2B from 0 to 18 h (= 3.64). The values of 7SDNA(0), *τ*, and *ϕ* were identical to those underlying the red 7SDNA(t) curve shown in Fig. 2A. By reducing the value of *ρ* for this curve by about 8% and using the same beta-binomial distribution to generate the initial dNTP pool size distribution as above (Fig. 2A), we obtained a WT N(t) curve that was acceptably similar to the target curve (Fig. 3A). In the two mutant cases, we used the same 7SDNA(t) curve and the same beta-binomial distribution as for the WT case. Furthermore, when creating the initial nucleoid population, we had to take into account that the dNTPs remaining after mtDNA replication are stored (*dNTP*_*rest*_) in daughter nucleoids and are later used for replication (see Fig. 3A caption for details). With this, we obtained N(t) curves for the two mutant lines that also were very similar to the target curve (Fig. 3A), while the cellular number of mtDNAs throughout the cell cycle (mtDNA(t)) were markedly different from the WT case (Fig. 3A).

**Figure 3.**
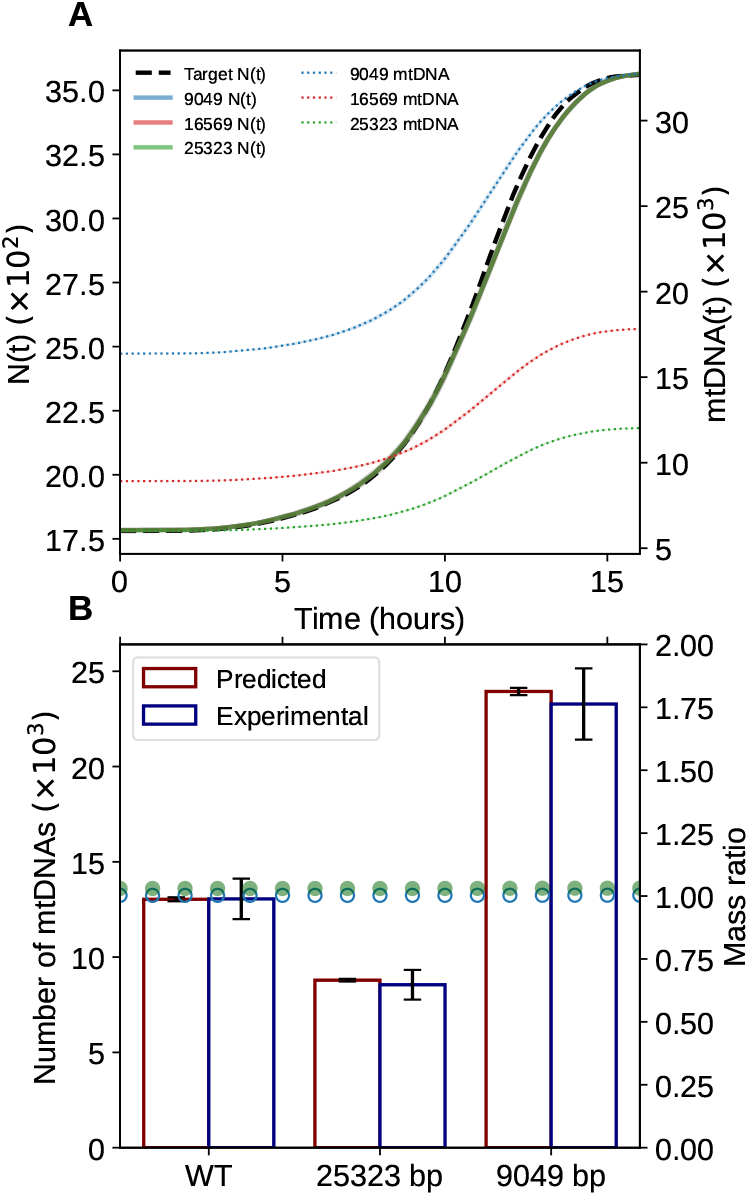
Nucleoid and mtDNA numbers in cybrids. A. Left y-axis: Mean number of nucleoids (N(t)) in the three cybrid lines (solid line) relative to the target N(t) profile. In all three cases the mean±SD number of nucleoids was 2,606±20.7. Parameter values used in Eq. (3): 7SDNA(0) = 1650 · 3.64, *τ* = 0.001, *ϕ* = 0.001, and *ρ* = 3.93. Parameter values used to generate the initial dNTP pool size distribution from a beta-binomial distribution: *α* = 10, *β* = 2. For each parameter combination, we used the mean N(t) from 50 samples from this distribution. In the 9049 bp case: 18% of the 1784 initial nucleoids contained 10 mtDNAs and their *dNT P*_*rest*_ values were randomly picked from the range [2808, 5614], while 82% contained 9 mtDNAs and their *dNT P*_*rest*_ values were randomly picked from the range [2808,18096]. In the 25323 bp case: 37% of the initial 1784 nucleoids contained 4 mtDNAs and their *dNT P*_*rest*_ values were randomly picked from the range [13752,27498], while 63% contained 3 mtDNAs and their *dNT P*_*rest*_ values were randomly picked from the range [13752,50640]. See https://github.com/Jagirhussan/mtDNA-mass-conservation for how these percentages and ranges were calculated. Right y-axis: Mean number of mtDNAs throughout the cell cycle in the three cybrid lines. B. Left y-axis: Comparison between estimated and reported (Tang et al., 2000) mean±SD number of mtDNA base pairs per cell when assaying a large population of asynchronous cells from the three cybrid lines. Right y-axis: Estimated mtDNA mass ratios between the 9,049 bp and 25,323 bp cybrid lines and the WT cybrid line (16,569 bp) throughout the cell cycle.

In the HeLa cell case, we predicted that the filling of the dNTP pools will be terminated in early G2 (Fig. S6). Assuming the existence of a generic regulatory scheme, this prediction is also likely to be applicable to 143B206 cells and can thus be used to test for consistency. In the 143B206 cell case, we predict that the dNTP pool size distribution will be similar to the initial distribution at t ≈ 17.5 hours (Fig. S9), which is consistent with the HeLa cell results.

In an ideal asynchronous cell culture, the probability distribution of where a cell is positioned on the time axis of the cell cycle will be approximately uniform. Hence, in a large sample of cells, the mean mtDNA number will approximately be the mean value of the mtDNA number throughout the cell cycle, and an mtDNA mass estimate (M) for each of the three cybrid lines can thus be obtained from the equation:

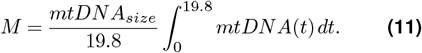

The mtDNA mass estimates corresponded very closely to the reported data (Tang et al., 2000) (Fig. 3B). Dividing the mtDNA mass for the 9,049 bp and 25323 bp cases with the WT mtDNA mass as a function of time shows that the cell cycle model predicts almost perfect mtDNA mass conservation throughout the cell cycle in both cases (Fig. 3B). This suggests that if one repeats the experiment reported by Sasaki *et al*. (Sasaki et al., 2017) in three cybrid lines and quantifies mtDNA numbers, we predict that almost perfect mtDNA mass conservation will be observed between the WT cybrid and the two others in all cell cycle phases. It should be noted that mass conservation and the fit between predicted and reported mtDNA numbers (Fig. 3B) are not dependent on the assumption that there are 5 mtDNAs per nucleoid in the WT line. If the mean number is lower or higher, this will only enforce a recalibration of the start and end point values of the N(t) curve (Fig. 3A).

The above results suggest that the increase in mtDNA number throughout the cell cycle in the three cybrid lines was based on the same regulatory scheme as in the HeLa cell case (Fig. 2), and indicates that our use of a simple single cell model to capture the average temporal dynamics of mtDNA number in a large population of cells is warranted.

## Discussion

The concept of an intranucleoid dNTP pool appears to provide a simple explanation for the observed conservation of the mtDNA mass in cybrid lines. And it provides the foundation for the notion that 7S DNA can be the main nucleotide provider for mtDNA replication in postmitotic, quiescent, and proliferating cells, offering at least one important proximate reason for the observation that 7S DNA has a very high turnover rate.

Although the cell cycle model is constrained well enough to make tentative conclusions, there is clearly a need for more comprehensive temporal and cell cycle stamped data from the same experimental system. This should include the number of nucleoids, the number of mtDNAs, the number of 7SDNAs and nucleoid replication time. If this in addition could be done with cybrid lines, one would obtain a data set on cellular dynamics that would allow a thorough scrutinization of the model’s premises and predictions, as well as provide guidance for experimental work targeting underlying molecular mechanisms. Furthermore, to our knowledge, there exists very little data on the concurrent temporal change in the number of mtDNAs, nucleoids, and 7SDNA in non-dividing cells. Such data would be instrumental for making progress in understanding how such cells can achieve homeostatic regulation of nucleoid and mtDNA numbers.

The observed mass conservation of mtDNA in cybrid lines (Tang et al., 2000) supports the notion that there must exist mechanisms that ensure the provision of a predefined number of dNTPs to each nucleoid that is based on a targeted number of wild-type mtDNAs. However, due to the paucity of relevant experimental data, it is premature to speculate on the regulatory architecture of the molecular machinery responsible for this dimensioning.

It was recently proposed that, in early passage non-cancerous adult human dermal fibroblasts (HDFa) obtained from healthy donors, only a minority of all nucleoids are actively involved in mtDNA replication (Brüser et al., 2021). This claim was based on the results of exposure of cells to 10 *µ*M of the thymidine analogue 5-ethynyl-2’-deoxyuridine (EdU) for 120 h. However, in Chinese hamster ovary (CHO) cells, cytotoxic effects already start to appear at a medium concentration of 0.05 *µ*M EdU when there is no natural thymidine in the medium (Ligasová et al., 2015; Haskins et al., 2020). A reason for this is that cells cannot easily use an EdU-containing strand as a template during DNA replication, possibly because the presence of EdU leads to replication fork stalling or collapse during DNA replication (Haskins et al., 2020). In the HDFa study 10% (v/v) fetal bovine serum was added to the medium, providing natural thymidine. Due to this, we think one should be cautious when interpreting data based on long-term EdU exposure. However, if it indeed turns out that only about 50% of nucleoids are involved in mtDNA replication during the cell cycle, this will not impact our assertions that the intranucleoid dNTP pool concept is of fundamental importance and that all dNTPs for mtDNA replication stem from 7SDNA produced within the nucleoid. The cell cycle model can easily be adapted to account for this possibility, but we deemed this unwarranted until more conclusive data emerge.

The production of dNTPs by the salvage pathway (Gandhi and Samuels, 2011) is energetically much less costly than the production of dNTPs by de novo synthesis (Wagner, 2005; Lynch and Marinov, 2015; Ghosh et al., 2023). The cost of adding two phosphate groups to a nucleoside monophosphate that comes from the degradation of 7S DNA is two ATPs (Blanco and Blanco, 2017), while the approximate cost of synthesis of a single dNTP molecule is estimated to be the hydrolysis of 50 ATP equivalents (Wagner, 2005; Lynch and Marinov, 2015; Ghosh et al., 2023). However, a thorough enzyme kinetic analysis showed that the rate of dNTP supply by the mitochondrial salvage pathway is so modest that even a single mtDNA in a mitochondrion will need more than 10 hours to replicate (Gandhi and Samuels, 2011). In our calculations on non-dividing cells above, the required number of dNTPs in the intranucleoid dNTP pool is, however, delivered over a period of 60-70 hours. Thus, the existence of individual nucleoid-associated dNTP pools would allow much more extensive use of the salvage pathway as a dNTP provider in these cells. In a standard reference person, muscle cells, adipocytes, and neurons represent more than 75% of the total cellular mass (Sender and Milo, 2021). If we count in quiescent mitotic cells, this suggests that in a predominant fraction of human cell mass, dNTPs that fuel mtDNA replication may come mainly from the salvage pathway. This figure is likely not very different in other mammals and in several other vertebrate species (Nicholls and Minczuk, 2014).

The above considerations suggest that the invention of an intranucleoid dNTP pool would have caused a fitness-impacting reduction in energy expenditure in non-dividing cells by drastically reducing the need for *de novo* synthesis of dNTPs to fuel mtDNA replication. A possible scenario is that the pool emerged due to the appearance of genetic variation that led to 7S DNA driven nucleotide production. Otherwise, an elaborate regulatory system would have had to be invented to control that the pool’s nucleotide composition closely matched the mtDNA nucleotide composition. The use of 7S DNA could also have been instrumental in bringing the number of nucleoids (and mtDNA) under tight dynamic cellular control. Irrespective of which selection force was the primary driver for bringing 7S DNA driven nucleotide production to the regulatory fore, our results indicate that one possible reason for evolutionary conservation of the D-loop in a wide range of species (Nicholls and Minczuk, 2014) is its key role in maintaining an intranucleoid dNTP pool.

## Acknowledgements

The authors wish to thank Goran Šimić and Daniel A. Beard for helpful comments on a previous version of this manuscript.

## Supplementary Information

**Figure S1.**
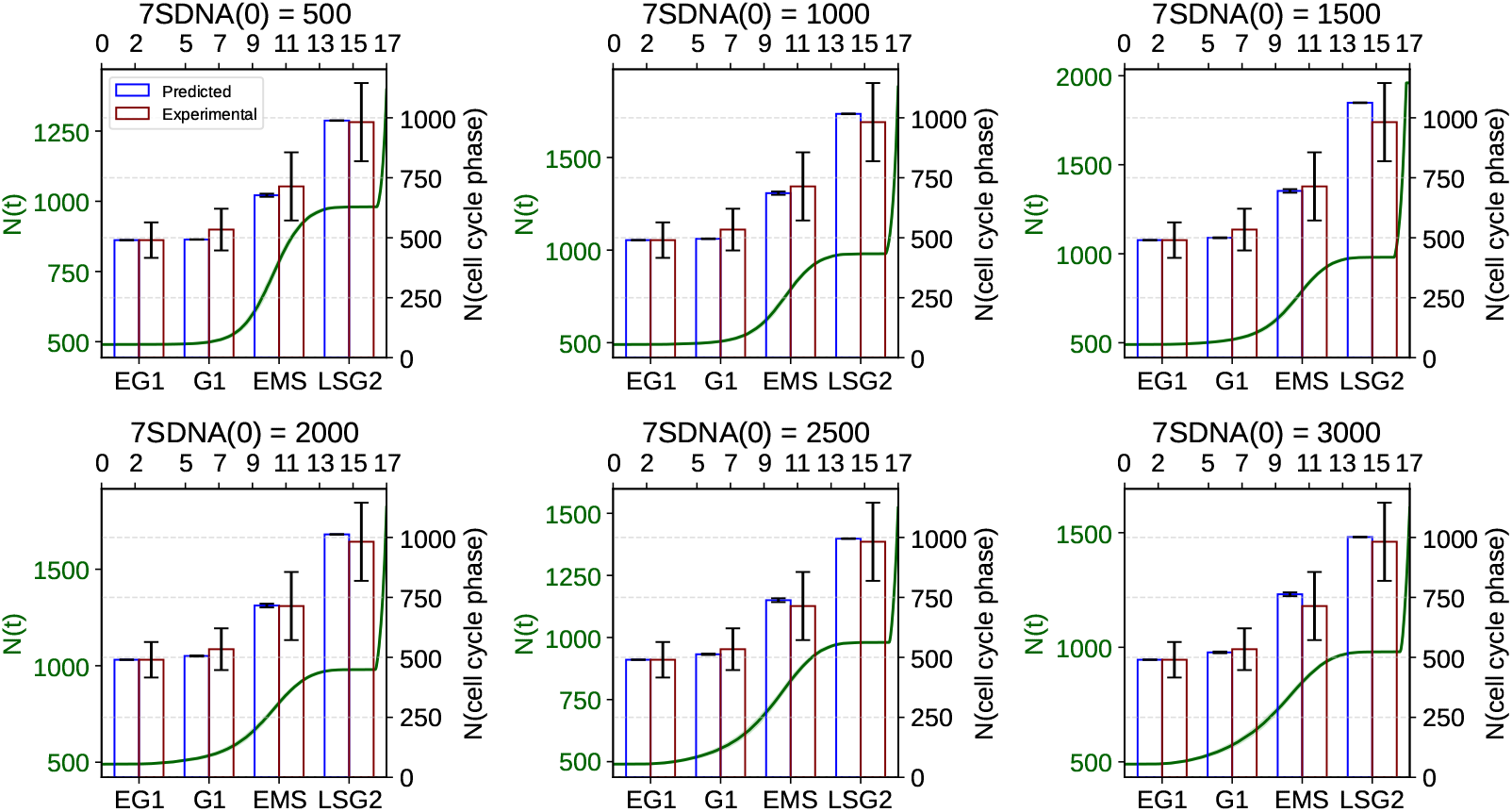
Top x-axis and left y-axis: Predicted number of nucleoids in a HeLa cell for six different 7SDNA(0) values from the beginning of EG1 (t = 0) to end of G2 (t = 17) when we let replication of nucleoids proceed unperturbed until end of G2. Each graph depicts the predicted mean ± SD trace obtained from 50 samples from the initial beta-binomial dNTP pool size distribution (*α*= 10, *β* = 2), but the variation is to small to be visible. Bottom x-axis and right y-axis: Predicted vs reported (Sasaki et al., 2017) number of nucleoids in a HeLa cell across its cell cycle for the same six 7SDNA(0) values. EG1, early G1 phase; EMS, early S/mid S phase; LSG2, late S/G2 phase.

**Figure S2.**
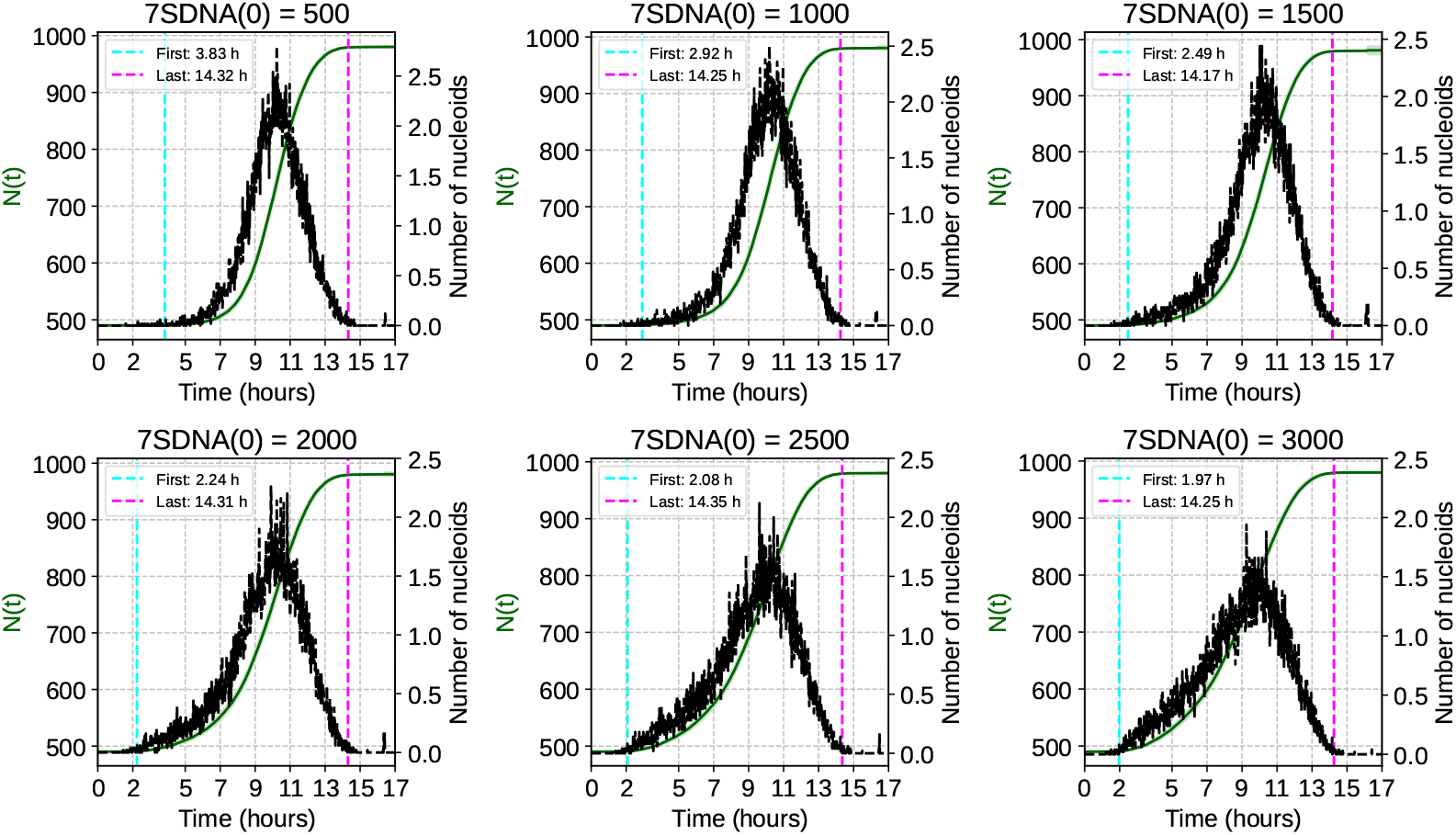
Left y-axis: The predicted number of nucleoids (N(t)) from the beginning of EG1 until end of G2 for the six 7SDNA(0) values used. Right y-axis: The number of nucleoid divisions in the same time frame. The vertical lines denote the time points of the first and last nucleoid division, respectively.

**Figure S3.**
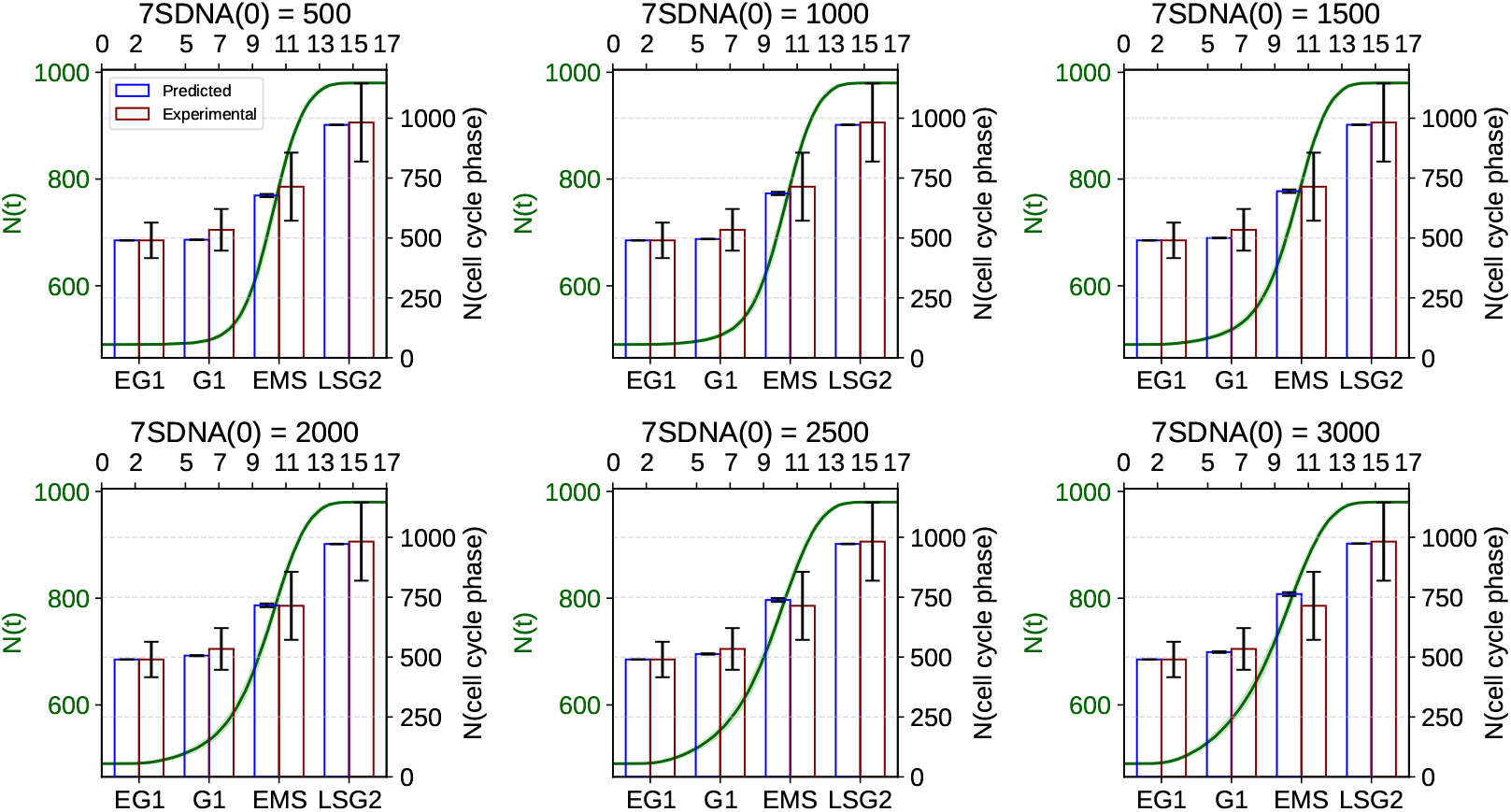
Top x-axis and left y-axis: Predicted number of nucleoids in a HeLa cell for six different 7SDNA(0) values from the beginning of EG1 (t = 0) to end of G2 (t = 17) when we assume that replication of nucleoids is inhibited shortly after duplication of the initial population of nucleoids. Each graph depicts the predicted mean ± SD trace obtained from 50 samples from the initial beta-binomial dNTP pool size distribution (*α* = 10, *β* = 2), but the variation is to small to be visible. Bottom x-axis and right y-axis: Predicted vs reported (Sasaki et al., 2017) number of nucleoids in a HeLa cell across its cell cycle for the same six 7SDNA(0) values when nucleoid production is inhibited shortly after duplication of the initial population of nucleoids. EG1, early G1 phase; EMS, early S/mid S phase; LSG2, late S/G2 phase.

**Figure S4.**
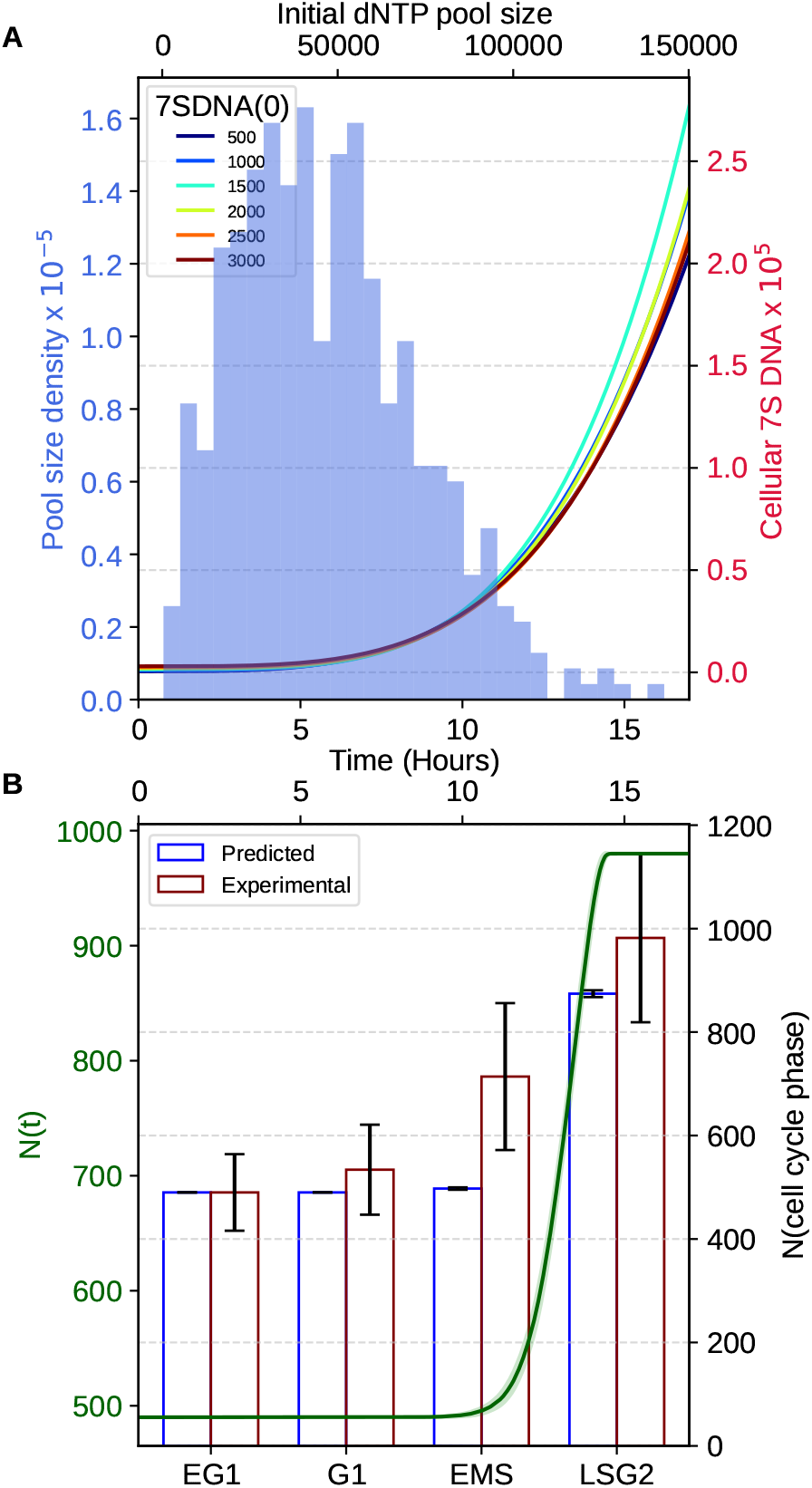
A. Top x-axis and left y-axis: Example of an initial dNTP pool size distribution based on a discrete beta-binomial distribution with *n* = 165690, *α* = 2 and *β* = 5. Bottom x-axis and right y-axis: Right y-axis: Estimated cellular number of 7S DNA throughout the cell cycle until end of G2 (t = 17) as a function of the initial cellular number of 7S DNA (7SDNA(0)) based on Eqs. (2) and (3). B. Top x-axis and left y-axis: Predicted number of nucleoids (N(t)) from the beginning of EG1 (t = 0) to end of G2 (t = 17) when we assume that replication of nucleoids is inhibited shortly after duplication of the initial population of nucleoids. The graph shows the mean±SD when using all the six 7SDNA(0) values depicted in panel A. Bottom x-axis and right y-axis: Predicted (mean±SD) vs reported (Sasaki et al., 2017) number of nucleoids (N(cell cycle phase)) in a HeLa cell across its cell cycle. EG1, early G1 phase; EMS, early S/mid S phase; LSG2, late S/G2 phase.

**Figure S5.**
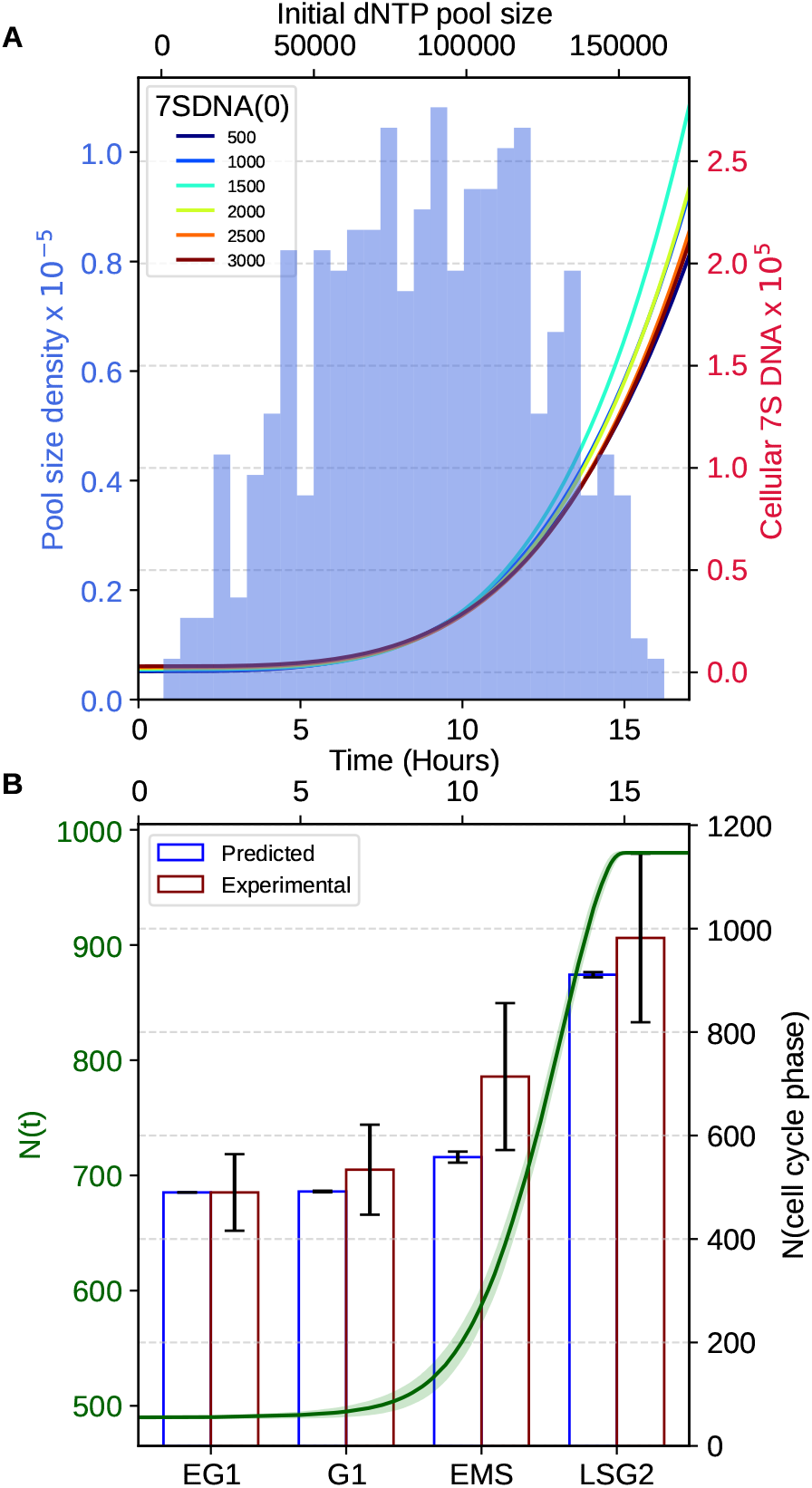
A. Top x-axis and left y-axis: Example of an initial dNTP pool size distribution based on a discrete beta-binomial distribution with *n* = 165690, *α* = 2 and *β* = 2. Bottom x-axis and right y-axis: Right y-axis: Estimated cellular number of 7S DNA throughout the cell cycle until end of G2 (t = 17) as a function of the initial cellular number of 7S DNA at t = 0 (7SDNA(0)) based on Eqs. (2) and (3). B. Top x-axis and left y-axis: Predicted number of nucleoids from the beginning of EG1 (t = 0) to end of G2 (t = 17) when we assume that replication of nucleoids is inhibited shortly after duplication of the initial population of nucleoids. The graph shows the mean±SD when using all the six 7SDNA(0) values depicted in panel A. Bottom x-axis and right y-axis: Predicted (mean±SD) vs reported (Sasaki et al., 2017) number of nucleoids in a HeLa cell across its cell cycle. EG1, early G1 phase; EMS, early S/mid S phase; LSG2, late S/G2 phase.

**Figure S6.**
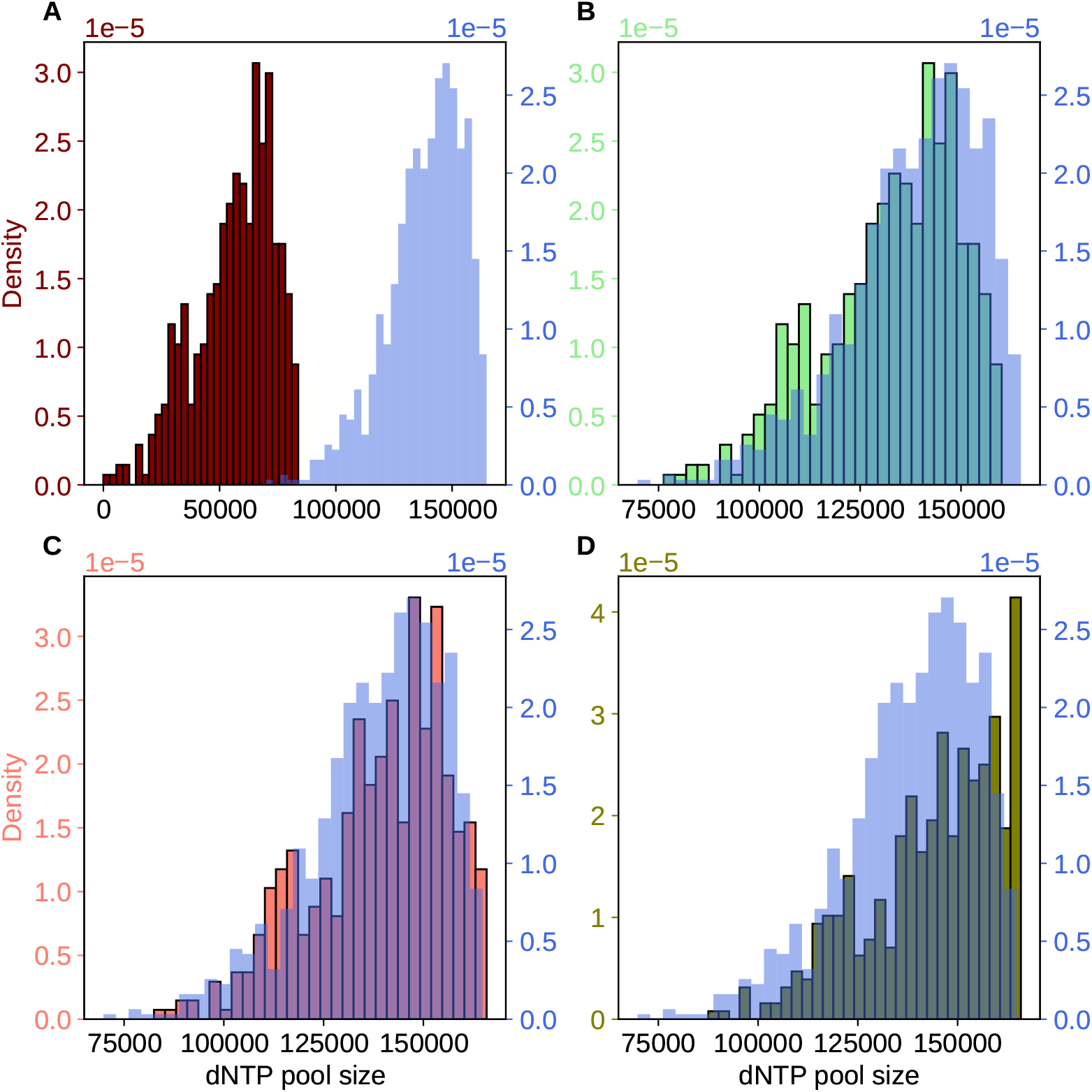
The dNTP pool size distribution vs initial distribution (light blue) at the time point when the initial population of nucleoids has duplicated and 7S DNA production is terminated(A), at t = 16.0 (beginning of G2) (B), at t = 16.5 (middle of G2) (C), and t = 17 (end of G2) (D).

**Figure S7.**
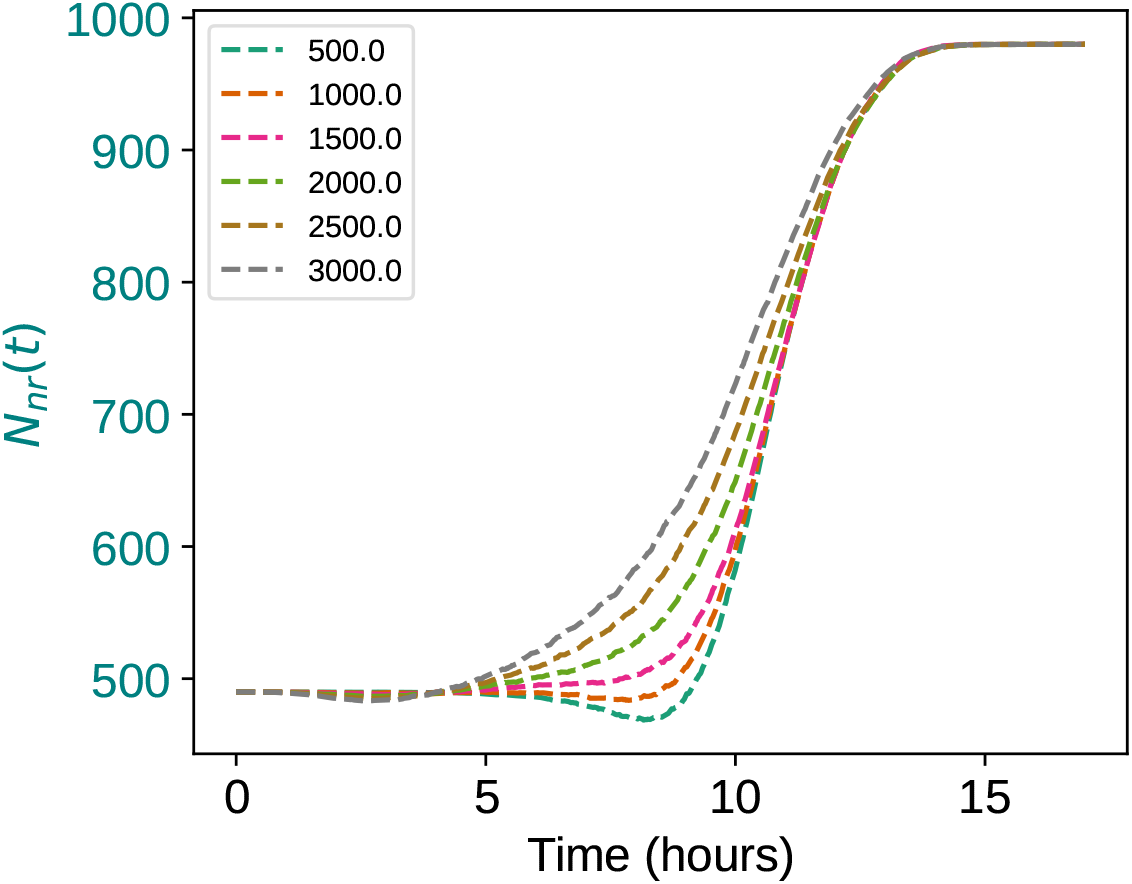
Predicted cellular number of nucleoids not actively replicating as a function of time (*N*_*nr*_(*t*)) for the six 7SDNA(0) values used.

**Figure S8.**
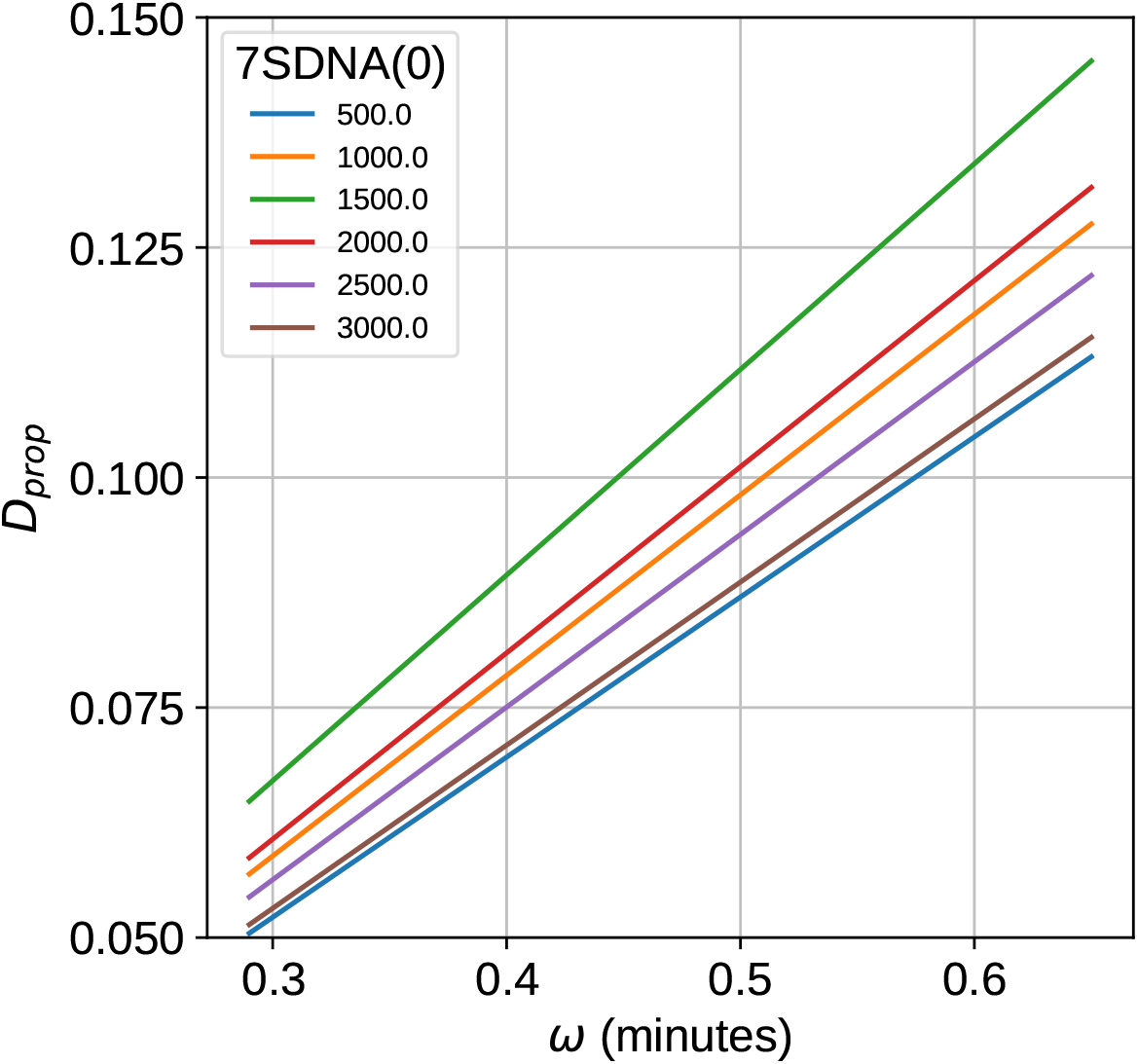
Predicted proportion of mtDNAs containing a D-loop as a function of variation in the 7S DNA synthesis time and the synthesis plus retention time (*ω*) and the initial cellular number of 7S DNA at t = 0 (7SDNA(0)).

**Figure S9.**
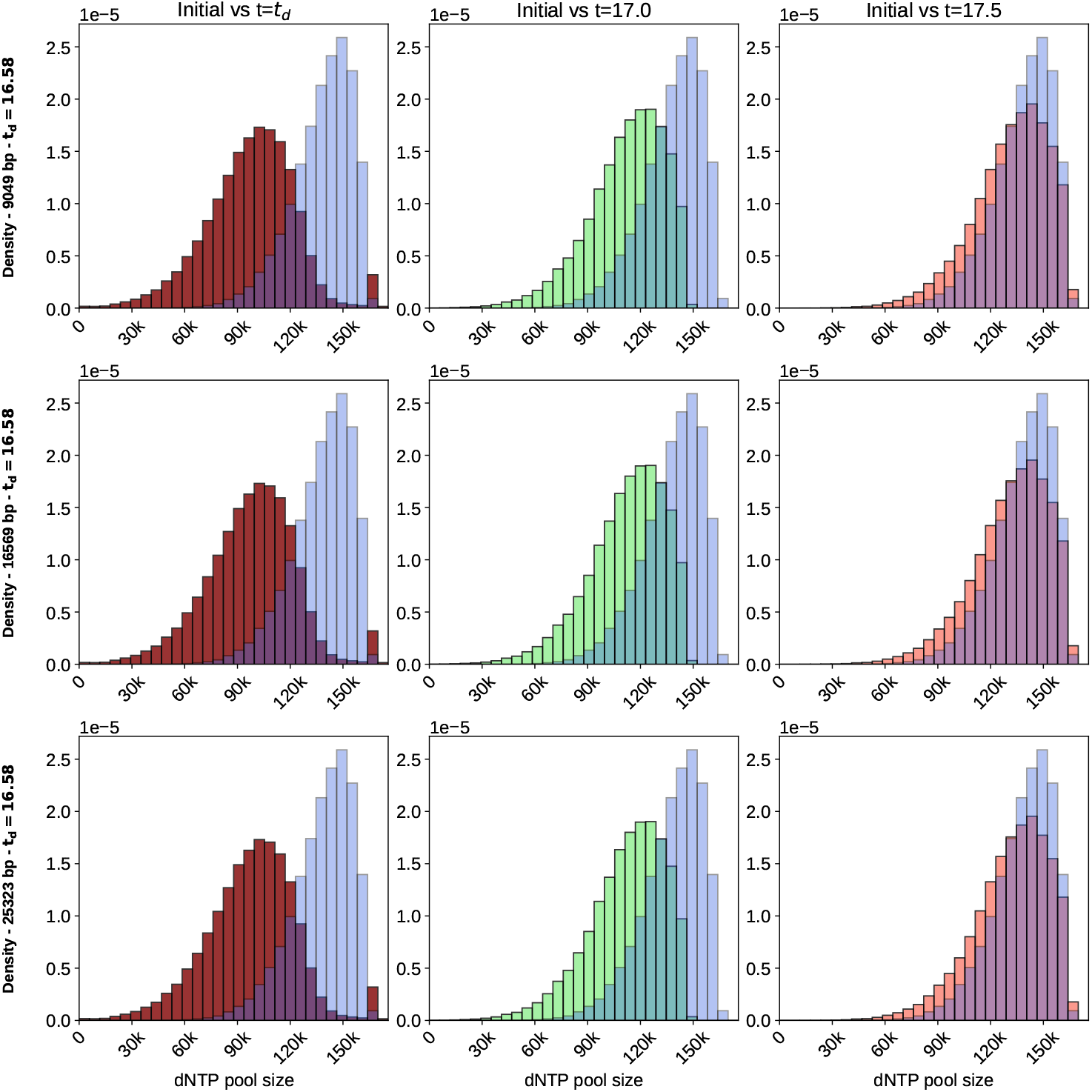
Comparison between the initial dNTP pool size distribution (light blue) vs the predicted distributions at t = *t*_*d*_, t = 17 and t = 18 in the tree cybrid 143B206 lines.

